# Dads on duty: First account of nest sitting in barnacle ganders

**DOI:** 10.1101/2024.07.19.604238

**Authors:** Isabella B. R. Scheiber, Annabel J. Slettenhaar, M. J. J. E. Loonen, Margje E. de Jong

## Abstract

In most Anseriformes (ducks, geese and swans) only females are known to incubate. Here we scientifically describe, indicidents of male nest sitting in barnacle geese (*Branta leucopsis*) as a form of paternal care of nest attendance. Based on pictures from wildlife cameras we identified males, which sat on their nests when their mates took incubation recesses. Wildlife cameras were placed at nests of which either the male or female was fitted with a GPS neck collar in the year prior, which aided with identifying individual birds on the nest. To attach transmitters, some geese were caught while defending their nests, thus we may have unintentionally selected bolder males which defended their nests more aggressively and were easier to catch. Nest sitting occurred relatively frequent, *i*.*e*. in 6/15 individuals. Our results show that males with collars were more likely to nest sit, but this does not deflect from the fact this behaviour exists in geese. In the course of this finding we discuss several possible functions of this behaviour, *i*.*e*. against raiding of nests by aerial predators, thermal control of nest temperature, and intraspecific brood parasitism. At this time we cannot demonstrate a possible function, as chances of successful hatching were not increased in nest-sitting males and we lack sample size for more in depth analyses. Lastly, we argue that ‘male incubation’ is misleading in the waterfowl literature, as it is truly justified for only two species, the black swan (*Cygnus atratus)* and black-bellied whistling duck (*Dendrocygna autumnalis)*.

## 1 INTRODUCTION

Nest defence by parents is considered a risky behaviour, as they may endure injury or even death when trying to ward off predators (de Jong, Nicolaus, Fokkema, & Loonen, 2021; Samelius & Alisauskas, 2006). Furthermore, nest defence also bears energetic costs (Caro, 2005; Thys, Lambreghts, Pinxten, & Eens, 2019) thus the risk of clutch protection must be gauged against its benefits (Ringelman & Stupaczuk, 2013). In birds, there is large variation in how much males and females contribute to parental care and in more than half of all species, males participate in incubation (Deeming & Reynolds, 2015). In most waterfowl species, *i*.*e*. swans, geese and ducks, however, only females incubate. For barnacle geese (*Branta leucopsis*) breeding on Svalbard anecdotal evidence indicates that males sometimes sit on the nest (from now on referred to as ‘nest sitting’). In this study we dive further into this behaviour.

The barnacle goose is an Arctic breeding species, renowned for nesting high on steep mountain cliffs and islets (Mitchell, Black, & Evans, 1998). They often breed on islands with glaucous gulls (*Larus hyperboreus*), Arctic skuas (*Stercorarius parasiticus*) and great skuas (*S. skua*), which are the main avian predators of barnacle geese eggs and/ or adults (Prop et al., 2015; Tombre & Erikstad, 1996). Similar to other geese (*e*.*g*. lesser snow goose, *A. caerulescens caerulescens;* pink-footed goose *A. brachyrhynchus;* Canada goose, *B. canadensis)*, barnacle geese protect their eggs against avian predators through a high rate of nest attendance (Prop, Van Eerden, & van Drent, 1984; Samelius & Alisauskas, 2001; Schreven, Stolz, Madsen, & Nolet, 2021) and active defence (Clermont, Réale, & Giroux, 2019; de Jong et al., 2021; Dittami, Kennedy, & Thomforde, 1979; Speelman, Hammers, Komdeur, & Loonen, 2022).

Just as in all other geese, only barnacle females are known to incubate. During nest construction, geese develop thermosensitive brood patches, which guarantee an efficient heat transfer from the body to the egg (Jones, 1971). During this time, ganders closely guard their receptive mates (Lamprecht, 1989). Males stay in close vicinity of the nest, maintaining contact with their incubating mates visually and vocally (Prop et al., 1984). In case of an intruder coming close, most males actively protect and defend the nest and incubating female (Speelman et al., 2022).

Incubation starts after the penultimate egg is laid (Hübner, Tombre, & Erikstad, 2002) and lasts for 25 days on average (Black, Prop, & Larsson, 2014). Females only take brief incubation recesses for feeding and preening, as each time they leave the nest, the eggs are at risk of dropping below an optimal temperature, possibly overheating, or being discovered by a predator (Ahmad & Li, 2023; Alsos, Elvebakk, & Gabrielsen, 1998). Before leaving the nest, females often use the down lining of the nest feathers to cover the eggs for insulation and/or hide them from predators. Common comprehension is that males either join their mates during incubation recesses, leaving the nest unattended, or stay behind close to the nest (Owen & Wells, 1979).

Here we report of several cases, in which some barnacle ganders nest sit, while females are away. First sporadic notes of this behaviour stem from 1979 in a barnacle goose colony on Nordenskiöldkysten (J. Prop pers. comm.) and 2006 in our Kongsfjorden study colony database (M. J. J. E. Loonen, pers. obs.). This type of behaviour was also mentioned anecdotally in a study on pink-footed geese breeding on Iceland, which described a few males ‘crouching over the eggs’ for up three minutes when their female left (Inglis, 1977). We decided to investigate this phenomenon further and provide the first scientific account of nest sitting in any goose species, although we cannot exclude the possibility that reports of this behaviour may have been disseminated outside peer-reviewed publications. We wanted to determine how often nest sitting of males occurred, if male age played a role, and whether individuals, which performed nest sitting did so multiple times or only once. Finally, we investigated if eggs were more likely to hatch if males were nest sitting.

## 2 METHODS

Our study population resides on the west coast of Spitsbergen, the largest island of the Svalbard archipelago. In Kongsfjorden an established breeding colony nests on a group of several islets (Tombre, Mehlum, & Loonen, 1998). These islets are characterised by exposed ridges and flat stretches of tundra (Tombre & Erikstad, 1996). Two are monitored in detail; the larger main breeding islet, Storholmen (ca. 30 ha, 259 nests in 2021) and Prins Heinrichøya (ca. 3 ha, 29 nests in 2021) located offshore of the village Ny-Ålesund (78°55’N, 11°56’E), Svalbard, Norway.

Within the scope of a larger study on circadian and circannual rhythmicity (de Jong et al., 2024) we fitted 24 adults (50% males and females of established pairs, respectively) in 2020 with solar-powered GPS-GSM transmitters attached with neckbands (OrniTrack-NL40 3G, Ornitela, UAB, Lithuania). Except for two geese, which were ringed as goslings, the individual age was back calculated from the time geese were originally colour marked as adults; thus, ages given in Table 1 are conservative. Individuals were chosen following a suite of criteria (for details see supplemental material) including that it would be desirable if both pair partners were marked with unique colour rings already. The neck collar allowed a definite identity recognition of pair partners on and close by the nest, even when colour rings were not visible. Sexing of all individuals was performed when birds were originally ringed by examining the cloaca for the presence of intromittent penises. To attach transmitters, geese were caught in the vicinity of the nest either by hand, with a small hand-net, or a fishing rod with an attached nylon snare (‘nest catches’, Fenstad et al., 2017), or else during the annually performed mass captures of moulting geese, where considerable numbers of geese are funnelled into a trap (‘moult catches’, Loonen, Oosterbeek, & Drent, 1997). Only males, which remained close to the nest when a human observer approached, could be caught by hand, using the fishing rod or hand-net. We are aware that these methods might comprise a bias in trappability *sensu* STRANGE, an acronym highlighting several possible sources of sampling biases (Webster & Rutz, 2020), as these males show high nest defence and risk-taking behaviour (de Jong et al., 2024). From here on we consider them ‘bold’.

**Table 1.** Identities of 15 focal pairs, whose nests were monitored with wildlife cameras in 2021. For each individual, we provide codes of unique colour (o=orange, g=green, y=yellow plus a three-letter code) and metal (CA***) rings, and if they were fitted with a transmitter, the neck collar ID. Except two individuals, which were ringed as goslings (*), the minimum age is given, calculated from the time the geese were originally ringed as adults (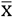 age ± SD: males 9.4 ± 3.31; females 10.4 ± 4.46). Whether or not males sat on nests is also indicated.

| Male colour ID | Male ID | Male neck collar ID | Age (years) | Male nest sitting | Female colour ID | Female ID | Female neck collar ID | Age (years) |
| --- | --- | --- | --- | --- | --- | --- | --- | --- |
| oFYJ | CA46944 | 201913 | 16 | yes | oFVD | CA46857 |  | 10 |
| gZUN | CA45605 | 201915 | 9 | yes | oFLC | CA45810 |  | 12 |
| alur | CA47072 | 201917 | 8 | no | yBSV | CA45997 |  | 4 |
| oFZY | CA46942 | 201918 | 10 | yes | gSNF | CA36931 |  | 19 |
| oFIJ | CA45800 | 201924 | 12 | yes | yBHJ | CA45891 |  | 5 |
| oFIA | CA45795 | 201930 | 12 | yes | gLDC | CA41238 |  | 14 |
| yBID | CA46840 | 201932 | 7 | no | yBIB | CA44228 |  | 14 |
| oFZX | CA46941 |  | 10 | no | gTTL | CA41113 | 201912 | 16* |
| yBYI | CA45746 |  | 7 | yes | yAUF | CA45744 | 201919 | 7 |
| yAXY | CA45703 |  | 8 | no | yBHY | CA46848 | 201925 | 5* |
| gLHX |  |  | 2 | no | yAVV | CA45736 | 201926 | 7 |
| yAZT | CA45715 |  | 8 | no | oFXV | CA45632 | 201927 | 10 |
| yAXC | CA45942 |  | 8 | no | yAPH | CA45929 | 201931 | 8 |
| gTCZ | CA46957 |  | 10 | no | yAXV | CA45701 | 201934 | 8 |
| yBYX | CA41397 |  | 14 | no | yBYV | CA41400 | 201935 | 14 |

In 2021, we were able to locate 15 nests of the initial 24 pairs, where either the male (n=7) or female (n=8) was fitted with a transmitter in the previous year. During transmitter attachment, four males were caught at the nest, while three were caught during the moult catch. In the following we refer to unique colour ring codes when mentioning certain individuals.

Near the 15 nests, we set up wildlife cameras (Usogood TC30 Trail Camera) to get a more detailed picture of behaviours of males and females during incubation. Cameras were used in the past and do not seem to disturb the geese (*e*.*g*. de Jong, Wetherbee, & Loonen, 2019). At each nest, we placed the camera, set in time-lapse mode, at a one-metre distance of the nest to monitor primarily the behaviour of the incubating female. Cameras took two pictures every five minutes to detect any movement of the geese. The time in which cameras provided reliable pictures varied between nests (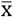 days ± SD: 12 ± 6.1; for details see supplement, Table S1, de Jong et al., 2024), thus we captured part of the incubation period. When scanning the photos, we noticed that in several instances males, rather than females, were sitting on the nest (Figure 1). As successive photos in a series are contingent, we only counted male nest sitting bouts as independent whenever the female returned and sat to the nest.

**Figure 1:**
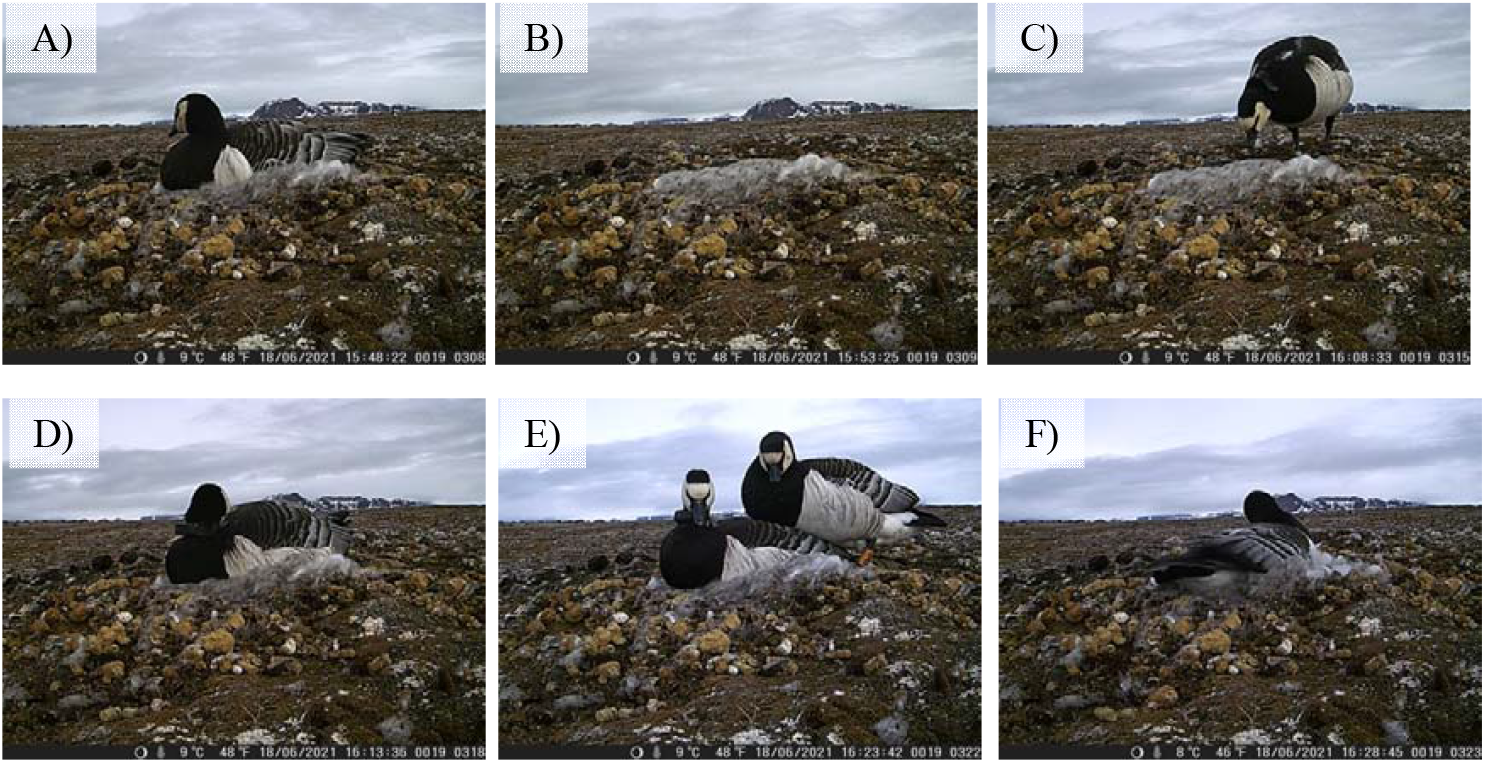
An example of one barnacle gander nest sitting. Photos taken on June 18th, 2021, between 15:48 and 16:28. Panel A) The female without a neck collar, fitted with the individual orange colour ring oFVD, is incubating. Panel B) The female left the nest. Panel C) Male alur with neck collar approaches the nest. Panel D) He sits on the nest with the neck collar clearly visible. Panel E) The female (with the colour ring partially visible) returns to the nest. Panel F) She resumes incubation. Temperature and time are shown in the lower right-hand corner of each photo.

### 2.1 Statistical Analysis

Using the R programming environment (R Core Team, 2021), we (1) applied a Student’s *t*-test to determine whether male age influenced the propensity to nest sit. We applied Fisher’s Exact Tests and give odds ratios and their 95% confidence intervals to determine, (2) if males, which sat on the nest, were more likely to have been fitted with transmitters, and (3) if male nest sitting was associated with hatching probability of the clutch, *i*.*e*. at least one egg hatched (for details on counts see 2 x 2 contingency tables, supplement Table S2). The sample size pertaining to the method of catching (nest catch vs. moult catch) was very small and showed clear separation, because all males that were caught at the nest showed nest sitting (Hauck & Donner, 1977). Due to the challenges this poses for statistical testing, we have opted to refrain from a formal statistical analysis in this case.

## 3 RESULTS

Observations of the 15 nests where cameras were placed revealed that six males sat on the nest, whereas nine did not (Table 1). Age of males had no effect on nest sitting [n _males not nest sitting_ = 9, 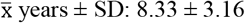; n _males nest sitting_ = 6, 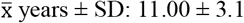; *t* _(13)_ = 1.612, p = 0.131, Figure 2a)]. Of the six nest sitting males, three were observed on the nest only once (gZUN, oFIJ, oFIA), whereas the others three males were observed multiple times (oFYJ and yBYI four times, oFZY six times). Males with neck collars were more likely to sit on nests than males without [Fisher’s Exact Test: p = 0.041, OR_2-sided_ = 13.59 with 95% CI of [0.86, 934.01], Figure 2b)]. Among males caught near the nest, 100% were observed sitting on nests (N = 4), compared to 33.3% of males caught during moult (N on nest = 1, N not on nest = 2, Figure 2c). Whether males nest sat had no effect on whether eggs hatched or else were abandoned/ depredated [Fisher’s Exact Test: p = 0.329, OR_2-sided_ = 0.3 with 95% CI of [0.02, 4.30], Figure 2d), see Table S2 for 2 x 2 contingency tables]. All but one male (oFIA), which started to nest sit shortly after the eggs hatched (see supplement Figure S1), sat on the nest during incubation. Males took on average 12 minutes (SD ± 17 min, range 0 – 42 min) after the female left before they started nest sitting and they averagely stayed for 39 minutes (SD ± 21 min, range = 12 – 70 min). This corresponds to about 60% (SD ± 27%, range = 16 – 88%) of the time the female was on incubation recess. After males terminated nest sitting, it took approximately 17 minutes (SD ± 13 min, range = 3 – 32 min) for the female to return to the nest (for details see Table S3). Based on examinations of four nests, where we had at least three days of camera observations (N = 4), the percentage of time when males nest sat during the females’ incubation recesses varied greatly between individuals (mean 20.0 % of the time, SD ± 21, range = 1.02 – 43%, for details see Table S4). However, throughout this study sample size is small, and these results should be taken with caution.

**Figure 2:**
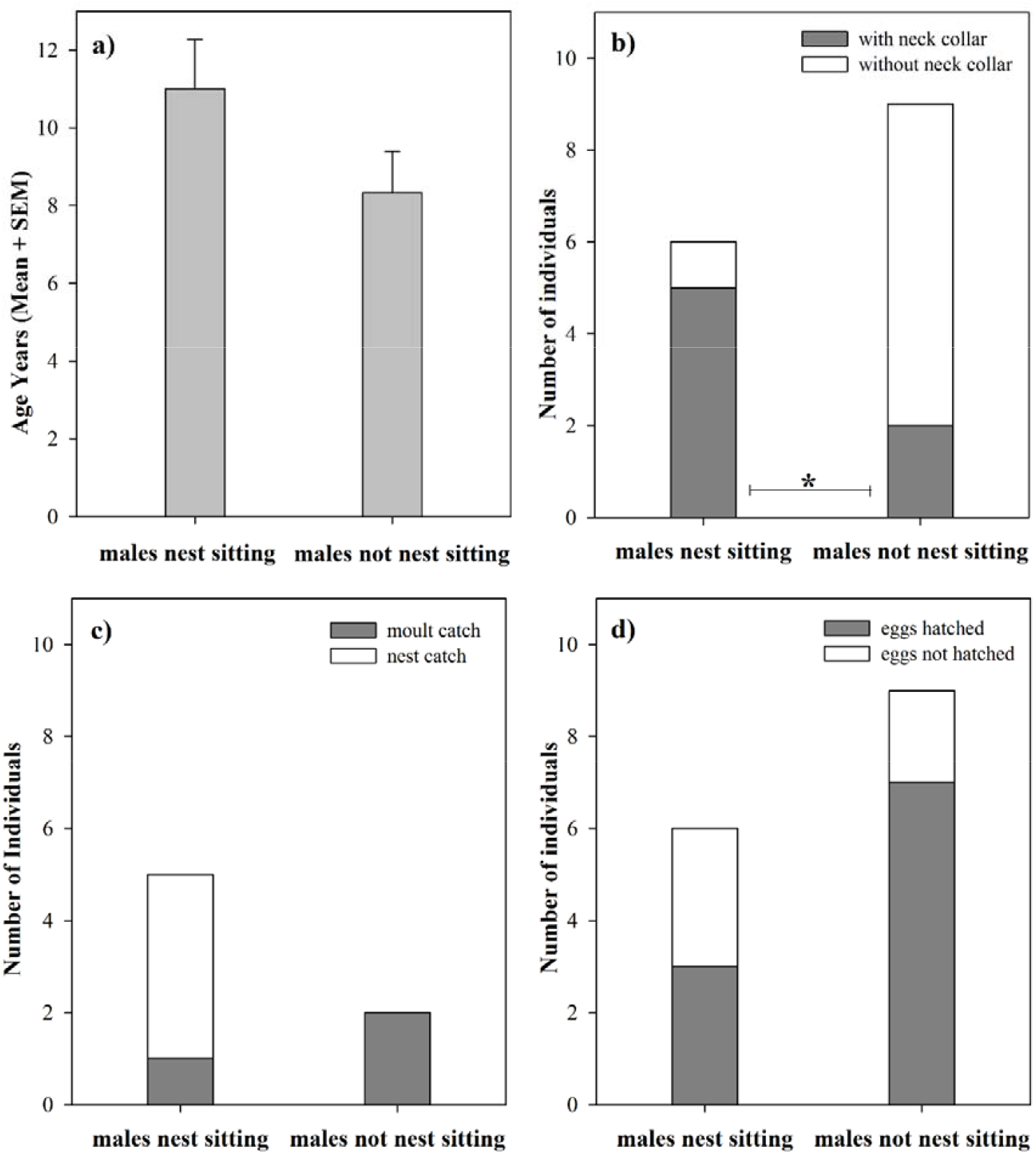
Features, which might influence nest sitting in barnacle ganders. a) Age did not affect the likelihood of male to nest sit (Student’s *t*-test: p > 0.05, whiskers represent the SEM). b) Males, which were fitted with a neck collar, were more likely to nest sit than males without (Fisher’s Exact Test: p = 0.041). c) Among males caught near the nest, 100% were observed to nest sit, while only 33,3% of males caught during moult did so. d) Whether eggs hatched or did not hatch was independent of males’ nest sitting (Fisher’s Exact Test: p > 0.05).

## 4 DISCUSSION AND CONCLUSION

### 4.1 Nest sitting in barnacle ganders – possible functions

To the best of our knowledge, this is the first scientific account of males sitting on nests in any goose species. Although we may have introduced a bias to catch predominantly bold individuals (Webster & Rutz, 2020), this does not alter the fact that some ganders perform this behaviour. We oppose to calling this behaviour ‘male incubation’, as termed repeatedly in the waterfowl literature (*e*.*g*. Brugger & Taborsky, 1994; Bruggers, 1979; Hawkins, 1986; Rollin, 1957), because there cannot be active transfer of heat from the male to the eggs as ganders do not form brood patches. Thus, we revive the term ‘nest sitting’ (Henson & Cooper, 1992) as the appropriate label. We identified from the photos that male posture on the nest was somewhat different from the female posture during incubation, *i*.*e*. males sat higher than females, not in the nest bowl but rather on top of the nest (compare panels A) and D) in Figure 1). Female ducks and geese sometimes adopt such as posture not only before actual incubation starts, but also during incubation (Hartman et al., 2023; Lorenz, 1991; Poussart, Larochelle, & Gauthier, 2000) to shield the nest and eggs from *e*.*g*. predation, inclement weather and/or intraspecific brood parasitism. We propose that male nest sitting has equivalent functions in the absence of the female. To date, we cannot provide a definite explanation of why this behaviour occurs, as neither the age of males seems to play a role, nor could we show an increase in hatching success of nests, where males nest sat. At this moment we lack a large enough sample for more in-depth analyses, such as when nest sitting occurs across the nesting period or to what extent males may compensate for more or longer incubation recesses of their mates. Furthermore, it should also be investigated in-depth, whether males adjust their behaviour in response to certain environmental conditions, such as nest density and location, and whether nest sitting confers fitness benefits (Madsen et al., 2019; Mery & Burns, 2010). In the following we suggest three possible, not mutually exclusive, functions of male nest sitting to initiate future studies not only in barnacle geese, but also other goose species.

The primary cause of hatching failure in ground nesting waterfowl is predation, thus various strategies are employed for nest protection (Peterson et al., 2022; Quinn, Prop, Kokorev, & Black, 2003; Samelius & Alisauskas, 2001; Schranck, 1972). Geese often nest near their main aerial predators. To not leave the nest unattended when the females are away, nest sitting of barnacle ganders might function in shielding the nest. This is backed by our choosing the most aggressive males when attaching transmitters, which might be bold enough to attack predators actively. In the future, attention should be paid to how often nest sitting of males occurs, how long the incubation recesses of the female last, what the predation pressure is, and at which stage during incubation the behaviour is performed to get a deeper understanding of male nesting as one form of predator protection.

Another possible function of nest sitting by males is to control the temperature in the nest. To some extent, nest insulation can reduce both the degree of egg cooling when the female is absent, and the time spent to rewarm the eggs after she returns from an incubation recess (Thompson & Raveling, 1988). Indeed, Arctic geese adjust their incubation behaviour in response to prevailing weather conditions (Elkins, 2004; Harvey, 1971; Poussart, Gauthier, & Jacques, 2001). Yet, egg cooling in the absence of the female might be overrated as even under unfavourable weather conditions, egg temperature drops during incubation recesses are nowhere near the minimal temperatures, which endanger an embryo’s development or survival (Poussart et al., 2000). It is worth noting that also overheating of unprotected eggs in the nest might become an issue in the Arctic in the future. Even if barnacle nests are exposed to direct sun, they probably will, at present, not reach this temperature limit long enough for embryos to die. Yet, common guillemots (*Uria aalge*) already showed an increased probability of egg loss at higher temperatures in a high-latitude colony in the Baltic Sea (Olin et al., 2024). Taken together, although nest sitting of barnacle ganders might assist in temperature control of eggs in the nest, we suggest it to only play a minor role, at least here and now.

If male nest sitting occurs during the period of egg laying, another possible function might be to prevent intraspecific brood parasitism, which frequently occurs in barnacle geese (Black et al., 2014). We cannot answer this question adequately, however, because we placed cameras not until the host female had started incubation, therefore lack photos from the laying period.

### 4.2 ‘Male incubation’ in waterfowl?

In the waterfowl literature only swans, and to some extent whistling ducks are described to deviate from the female only incubation pattern in waterfowl, as here male incubation occurs (Brugger & Taborsky, 1994; Hawkins, 1986; Scott, 1977). But are these males indeed incubating or is it a term that should be avoided, as is leads to a misconception when compared with species, where incubation is truly shared (Deeming & Reynolds, 2015)?

Hawkins already concluded that male incubation in tundra swans (*Cygnus columbianus columbianus*.) was not essential for successful embryo development (Hawkins, 1986) but protected against egg predation and provided some egg cooling benefits. This led Henson & Cooper to apply the term ‘nest sitting’ in trumpeter swans *(C. buccinator*), as here males occasionally exhibiting some of the nest-settling motions characteristic of incubating females, independent of predator presence or adverse weather conditions (Henson & Cooper, 1992). Our extensive search of the literature revealed that in most cases, in which male incubation was described (*e*.*g*. Bruggers, 1979; Flickinger, 1975; Rollin, 1957), they nest sit rather than incubate.

The two notable exceptions, for which the term incubation is warranted, are the black swan (*C. atratus*, BLSW, Cygnidae) and black-bellied whistling duck (*D. autumnalis*, BBWD, Anatidae). Whereas BLSW exhibit the typical long-term monogamy and biparental care of swans, this is true also for the BBWD, a feature atypical for ducks. Some other features, which the two species share are that neither females nor males develop brood patches (Bolen & Smith, 1979; Skutch, 1976), thus either parent is equally fit to incubate, and that both species may re-nest after successful breeding attempts (Bolen & Smith, 1979; Brugger & Taborsky, 1994; Coleman, 2014; Delnicki, 1973; James, Thompson, & Ballard, 2012). Most other waterfowl attempt a second clutch only if the first one failed. BLSWs and BBWDs might be able to re-nest because both parents incubate. Near constant incubation minimizes the time of egg development and hatching, thereby shortening the inter-clutch interval (James et al., 2012).

### 4.3 Conclusion

To sum up, this is the first scientific report in any goose species, where males expand paternal duties beyond the established view, namely that during the breeding period they only protect the nest and female in close vicinity of the nest. We have pictorial evidence that several barnacle ganders nest sat in the absence of their mates. Both predation and thermal risks are increased for eggs, which are not protected by parents, thus minimising those risks are the main causes for successful hatching. In barnacle geese, we suggest nest sitting of males to be a response to avian predator presence. We cannot exclude that preventing egg cooling in the absence of the female may also play a role but consider this unlikely. Our final point is that in most waterfowl species, males do not participate in incubation. Nest sitting is an unbiased term and should be used for cases, in which no active warming of eggs has (yet) been demonstrated.

## Data Availability Statement

The data that support the findings of this study are provided within the article and corresponding supplemental material.

## Acknowledgements

We are grateful to Børge Moe (Norwegian Institute for Nature Research, NINA) as the project administrator of AJS’ field grant, the staff of the French - German Arctic Research Base at Ny-Ålesund, chiefly Bettina Haupt (station leader AWI 2020/21), and KingsBay AS, Ny-Ålesund for logistic support. We thank Jouke Prop for sharing his unpublished observations and Kees Schreven for directing out attention to the one dated reference, where nest sitting in ganders was briefly alluded to. Comments by two anonymous reviewers and Thomas Lameris from the editorial board of *Ardea* helped to improve an earlier version of this manuscript. Funding was provided by the Austrian Science Fund (P32216 to I. B. R. Scheiber) and an Arctic Field Grant (The Research Council of Norway) in collaboration with NINA (ES676286 - to A. J. Slettenhaar). The study conforms to Directive 2010/63/EU and was conducted under FOTS ID 23358 from the Norwegian Animal Research Authority and approved by the Governor of Svalbard (RIS ID 11237).

## REFERENCES

Ahmad, I. M., & Li, D. (2023). More than a simple egg: Underlying mechanisms of cold tolerance in avian embryos. Avian Research, 14, 100104. doi: 10.1016/j.avrs.2023.100104

Alsos, I. G., Elvebakk, A., & Gabrielsen, G. W. (1998). Vegetation exploitation by barnacle geese Branta leucopsis during incubation on Svalbard. Polar Research, 17, 1–14. doi: 10.3402/polar.v17i1.6603

Black, J. M., Prop, J., & Larsson, K. (2014). The Barnacle Goose. London, UK: Bloomsbury Publishing.

Bolen, E. G., & Smith, E. N. (1979). Notes on the incubation behavior of black-bellied whistling ducks. Prairie Naturalist, 11(4), 119–123.

Brugger, C., & Taborsky, M. (1994). Male incubation and its effect on reproductive success in the black swan, Cygnus atratus. Ethology, 96(2), 138–146. doi: 10.1111/j.1439-0310.1994.tb00889.x

Bruggers, R. L. (1979). Nesting patterns of captive mandarin ducks. Wildfowl, 30, 45–54.

Caro, T. M. (2005). Antipredator Defenses in Birds and Mammals. Chicago, USA: University of Chicago Press.

Clermont, J., Réale, D., & Giroux, J.-F. (2019). Similarity in nest defense intensity in Canada goose pairs. Behavioral Ecology and Sociobiology, 73(8). doi: 10.1007/s00265-019-2719-3

Coleman, J. T. (2014). Breeding biology of the black swan Cygnus atratus in southeast Queensland, Australia. Wildfowl, 64, 217–230.

de Jong, M. E., Nicolaus, M., Fokkema, R. W., & Loonen, M. J. J. E. (2021). State dependence explains individual variation in nest defence behaviour in a long-lived bird. Journal of Animal Ecology, 90(4), 809–819. doi: 10.1111/1365-2656.13411

de Jong, M. E., Slettenhaar, A. J., Fokkema, R. W., Leh, M., Barnreiter, E., Griffin, L., … Scheiber, I. B. R. (2024). Diel and seasonal rhythmicity in activity and corticosterone in an Arctic migratory herbivore: A multifaceted approach. preprint. doi: 10.1101/2024.08.30.610510

de Jong, M. E., Wetherbee, R., & Loonen, M. J. J. E. (2019). Effects of fleas on nest success of Arctic barnacle geese: Experimentally testing the mechanism. Journal of Avian Biology, 50(5), e01944. doi: 10.1111/jav.01944

Deeming, D. C., & Reynolds, S. J. (2015). Nests, Eggs, and Incubation. Oxford, UK: Oxford University Press.

Delnicki, D. E. (1973). Renesting, incubation behavior, and compound clutches of the black-bellied tree duck in southern Texas. MSc, Texas Tech University, Lubbock, TX, USA. (31295001778181)

Dittami, J. P., Kennedy, S., & Thomforde, C. (1979). Observations on barnacle goose breeding, Branta leucopsis, in Spitsbergen 1975. Journal of Ornithology, 120, 188–195. doi: 10.1007/BF01642997

Elkins, N. (2004). Breeding: Incubation. In N. Elkins (Ed.), Weather and Bird Behaviour (Vol. 3rd edition, pp. 101–104). London, UK: T.& A.D. Poyser.

Fenstad, A. A., Bustnes, J., Lierhagen, S., Gabrielsen, K. M., Öst, M., Jaatinen, K., … Krøkje, Å. (2017). Blood and feather concentrations of toxic elements in a Baltic and an Arctic seabird population. Marine Pollution, 114(2), 1152–1158. doi: j.marpolbul.2016.10.034

Flickinger, E. L. (1975). Incubation by a male fulvous tree duck. Wilson Bulletin, 87(1), 106–107.

Hartman, C. A., Ackerman, J. T., Peterson, S. H., Fettig, B., Casazza, M., & Herzog, M. P. (2023). Nest attendance, incubation constancy, and onset of incubation in dabbling ducks. Plos One, 18(5), e0286151. doi: 10.1371/journal.pone.0286151

Harvey, J. M. (1971). Factors affecting blue goose nesting success. Canadian Journal of Zoology, 49(2), 223–234. doi: 10.1139/z71-03

Hauck, W. W., & Donner, A. (1977). Wald’s test as applied to hypotheses in logit analysis. Journal of the American Statistical Association, 72(360a), 851-853. doi: 10.1080/01621459.1977.10479969

Hawkins, L. L. (1986). Nesting behaviour of male and female whistling swans and implications of male incubation. Wildfowl, 37, 5–27.

Henson, P., & Cooper, J. A. (1992). Division of labour in breeding trumpeter swans Cygnus buccinator. Wildfowl, 43, 40–48.

Hübner, C. E., Tombre, I. M., & Erikstad, K. E. (2002). Adaptive aspects of intraclutch egg-size variation in the high Arctic barnacle goose (Branta leucopsis). Canadian Journal of Zoology, 80, 1180–1188. doi: 10.1139/z02-100

Inglis, I. R. (1977). The breeding behaviour of the pink-footed goose: Behavioural correlates of nesting success. Animal Behaviour, 25, 747–764. doi: 10.1016/0003-3472(77)90125-7

James, J. D., Thompson, J. E., & Ballard, B. M. (2012). Evidence of double brooding by black-bellied whistling-ducks. Wilson Journal of Ornithology, 124(1), 183–185. doi: 10.1676/11-082.1

Jones, R. E. (1971). The incubation patch of birds. Biological Reviews, 46(3), 315–339. doi: 10.1111/j.1469-185X.1971.tb01048.x

Lamprecht, J. (1989). Mate guarding in geese: Awaiting female receptivity, protection of paternity or support of female feeding. In A. E. Rasa, C. Vogel & E. Voland (Eds.), The Sociobiology of Sexual and Reproductive Strategies. Dordrecht, NL: Springer Science and Business Media.

Loonen, M. J. J. E., Oosterbeek, K., & Drent, R. H. (1997). Variation in growth of young and adult size barnacle geese Branta leucopsis: Evidence for density dependence. Ardea, 85, 177–192.

Lorenz, K. (1991). Here I Am - Where Are You? New York, NY, USA: Hartcourt Brace Jovanovich.

Madsen, J., Jaspers, C., Frikke, J., Gundersen, O. M., Nolet, B. A., Nolet, K., … de Vries, P. P. (2019). A gloomy future for light-bellied brent geese in Tusenøyane, Svalbard, under a changing predator regime. Polar Research, 38, 3393. doi: 10.33265/polar.v38.3393

Mery, F., & Burns, J. G. (2010). Behavioural plasticity: An interaction between evolution and experience. Evolutionary Ecology, 24(3), 571–583. doi: 10.1007/s10682-009-9336-y

Mitchell, C., Black, J. M., & Evans, M. (1998). Breeding success of cliff-nesting and island-nesting barnacle geese in Svalbard. Norsk Polarinstitutt Skrifter, 200, 141–146.

Olin, A. B., Dück, L., Berglund, P. A., Karlsson, E., Bohm, M., Olsson, O., & Hentati-Sundberg, J. (2024). Breeding failures and reduced nest attendance in response to heat stress in a high-latitude seabird. Marine Ecology Progress Series, 737, 147–160. doi: 10.3354/meps14244

Owen, M., & Wells, R. L. (1979). Territorial behaviour in breeding geese -A re-examination of Ryder’s hypothesis. Wildfowl, 30, 20–26.

Peterson, S. H., Ackerman, J. T., Keating, M. P., Schacter, C. R., Hartman, C. A., Casazza, M. L., & Herzog, M. P. (2022). Predator movements in relation to habitat features reveal vulnerability of duck nests to predation. Ecology and Evolution, 12(9). doi: 10.1002/ece3.9329

Poussart, C., Gauthier, G., & Jacques, L. (2001). Incubation behaviour of greater snow geese in relation to weather conditions. Canadian Journal of Zoology, 79, 671–678. doi: 10.1139/z01-023

Poussart, C., Larochelle, J., & Gauthier, G. (2000). The thermal regime of eggs during laying and incubation in greater snow geese. Condor, 102(2), 292–300. doi: 10.1093/condor/102.2.292

Prop, J., Aars, J., Bårdson, B.-J., Hanssen, S. A., Bech, C., Bourgeon, S., … Moe, B. (2015). Climate change and the increasing impact of polar bears on bird populations. Frontier in Ecology and Evolution, 3, 33. doi: 10.3389/fevo.2015.00033

Prop, J., Van Eerden, M. R., & van Drent, R. H. (1984). Reproductive success in the barnacle goose Branta leucopsis in relation to food exploitation on the breeding grounds, western Spitsbergen. Norsk Polarinstitutt Skrifter, 181, 87–117.

Quinn, J. L., Prop, J., Kokorev, Y., & Black, J. M. (2003). Predator protection or similar habitat selection in red-breasted goose nesting associations: Extremes along a continuum. Animal Behaviour, 65(2), 297–307. doi: 10.1006/anbe.2003.2063

R Core Team. (2021). R: A language and environment for statistical computing (Version 1.13). Vienna, Austria: R Foundation for Statistical Computing. Retrieved from https://www.R-project.org/.

Ringelman, K. M., & Stupaczuk, M. J. (2013). Dabbling ducks increase nest defense after partial clutch loss. Condor, 115(2), 290–297. doi: 10.1525/cond.2013.120096

Rollin, N. (1957). Incubation by drake wood duck in eclipse plumage. Condor, 59(4), 263–265. doi: 10.2307/1364656

Samelius, G., & Alisauskas, R. T. (2001). Deterring arctic fox predation: The role of parental nest attendance by lesser snow geese. Canadian Journal of Zoology, 79, 861–866. doi: 10.1139/z01-048

Samelius, G., & Alisauskas, R. T. (2006). Sex-biased costs in nest defence behaviours by lesser snow geese (Chen caerulescens): Consequences of parental roles. Behavioral Ecology and Sociobiology, 59, 805–810.

Schranck, B. W. (1972). Waterfowl nest cover and some predation relationships. The Journal of Wildlife Management, 36(1), 182. doi: 10.2307/3799210

Schreven, K. H. T., Stolz, C., Madsen, J., & Nolet, B. A. (2021). Nesting attempts and success of Arctic-breeding geese can be derived with high precision from accelerometry and GPS-tracking. Animal Biotelemetry, 9(1), 25. doi: 10.1186/s40317-021-00249-9

Scott, D. (1977). Breeding behaviour of wild whistling swans. Wildfowl, 28, 101–106.

Skutch, A. F. (1976). Parent Birds And Their Young: University of Texas Press.

Speelman, F. J. D., Hammers, M., Komdeur, J., & Loonen, M. J. J. E. (2022). Nest defence behaviour is similar between pair members but only male behaviour predicts nest survival in barnacle geese. Journal of Avian Biology, 2022(9), e02982. doi: 10.1111/jav.02982

Thompson, S. C., & Raveling, D. G. (1988). Nest insulation and incubation constancy of Arctic geese. Wildfowl, 39, 124–132.

Thys, B., Lambreghts, Y., Pinxten, R., & Eens, M. (2019). Nest defence behavioural reaction norms: testing life-history and parental investment theory predictions. R Soc Open Sci, 6(4), 182180. doi: 10.1098/rsos.182180

Tombre, I. M., & Erikstad, E. (1996). An experimental study of incubation effort in high-Arctic barnacle geese. Journal of Animal Ecology, 65, 325–331. doi: 10.2307/5878

Tombre, I. M., Mehlum, F., & Loonen, M. J. J. E. (1998). The Kongsfjorden colony of barnacle geese: Nest distribution and the use of breeding islands 1980-1997. Norsk Polarinstitutt Skrifter, 200, 57–65.

Webster, M. M., & Rutz, C. (2020). How STRANGE are your study animals? Nature, 582(7812), 337–340. doi: 10.1038/d41586-020-01751-5

