## Supplement for "Dads on duty: First account of nest sitting in barnacle ganders"

**Criteria for attachment of transmitters to selected focal individuals:**

1) Minimum age of three years
2) Both pair partners with uniquely coded colour rings desirable
3) Observed in Ny-Ålesund in two consecutive years to ensure that they went through at least one complete annual cycle
4) Preferably with known nest site locations
5) Found in and/or close to the village after hatching of young
6) Good health as judged by their physique and plumage condition

**Table S1:** Details of wildlife camera placement from 15 barnacle goose nests in 2021. For each individual we provide codes of unique colour rings, GPS transmitter ID and sex. Abbreviations of nest locations on islets in Kongsfjorden (see also Figure 1): SH = Storholmen, PH = Prins Heinrichøya, JH = Juttaholmen, OH = Observasjonsholmen, MH = Midtholmen. Date and time when cameras were placed and the number of days, when they, provided reliable pictures of observations of males and females on the nest, are provided. Nest fate indicates whether eggs hatched (H), nests were abandoned (A) or depredated (P).

| Nest Nr. | Goose Colour ID (Transmitter Nr.) | Sex | Nest  Location | Camera placed (Date, Time) | # days with at least one observation | Nest fate |
| --- | --- | --- | --- | --- | --- | --- |
| 1 | oFYJ (201913) | M | SH | 06/18, 02:55 | 3 | A/P? |
| 2 | gZUN (201915) | M | JH | 06/25, 14:04 | 1 | P |
| 3 | alur (201917) | M | SH | 06/18, 03:05 | 14 | H |
| 4 | oFZY (201918) | M | SH | 06/18, 01:48 | 4 | A |
| 5 | oFIJ (201924) | M | SH | 06/18, 01:17 | 15 | H |
| 6 | oFIA (201930) | M | SH | 06/18, 02:10 | 19 | H |
| 7 | yBID (201932) | M | SH | 06/18, 02:17 | 18 | H |
| 8 | gTTL (201912) | F | SH | 06/18, 03:22 | 15 | H |
| 9 | yAUF (201919) | F | SH | 06/18, 02:30 | 13 | H |
| 10 | yBHY (201925) | F | SH | 06/18, 02:41 | 16 | H |
| 11 | yAVV (201926) | F | MH | 06/25, 15:26 | 12 | H |
| 12 | oFXV (201927) | F | OH | 06/25, 14:52 | 14 | H |
| 13 | yAPH (201931) | F | JH | 06/25, 14:14 | 3 | P |
| 14 | yAXV (201934) | F | PH | 06/09, 15:22 | 15 | A/P? |
| 15 | yBYV (201935) | F | SH | 06/18, 01:05 | 18 | H |

**Table S2:** 2 x 2 contingency tables of males, which were or were not sitting on nests and if or if not they were a) fitted with neck collars; b) caught during moult or nest catches; and c) if eggs hatched or not.

|  |  | **Male sitting on nest** | **Male not sitting on nest** | ***Σ*** |
| --- | --- | --- | --- | --- |
| **a)** | **Male with neck collar** | 5 | 2 | ***7*** |
|  | **Male without neck collar** | 1 | 7 | ***8*** |
|  | ***Σ*** | ***6*** | ***9*** | ***15*** |
| **b)** | **Moult catch** | 1 | 2 | 3 |
|  | **Nest catch** | 4 | 0 | 4 |
|  | ***Σ*** | ***5*** | ***2*** | ***7*** |
| **c)** | **Eggs hatched** | 3 | 7 | 10 |
|  | **Eggs did not hatch** | 3 | 2 | 5 |
|  | ***Σ*** | ***6*** | ***9*** | ***15*** |

**Table S3:** Individual observations of male nest sitting behaviour. The duration of time when males were on the nest, then length of the female recess bout, the percentage of time the male spent on the nest in the absence of the female and the duration of time until the male nest sat after the female left as well as the duration of time it took the female to resume incubation after the male left are provided. For males, which nest sat multiple times, also individual means are provided (in italics). *Observation for one male, oFIA, was excluded, because he sat on the nest only after eggs hatched. His mate, gLDC, stayed in the vicinity of the nest with the newly hatched goslings, returning to the nest repeatedly (see Figure S1).

|  | **Duration male on the nest** | **Female recess bout length** | **% of time male nest sitting** | **Duration from female leaving to start male nest sitting** | **Duration from male leaving until female returns** |
| --- | --- | --- | --- | --- | --- |
| oFYJ | 00:15:09 | 00:35:21 | 42.86 | 00:20:12 | 00:00:00 |
| oFYJ | 01:00:34 | 01:00:34 | 100 | 00:00:00 | 00:00:00 |
| oFYJ | 00:10:06 | 02:21:19 | 7.15 | 00:00:00 | 02:11:13 |
| oFYJ | 01:10:40 | 01:15:43 | 93.33 | 00:05:03 | 00:00:00 |
| *Mean (oFYJ)* | *00:39:07* | *01:18:14* | *60.84* | *00:06:19* | *00:32:48* |
| yBYI | 00:05:04 | 00:30:17 | 16.73 | 00:10:05 | 00:15:08 |
| yBYI | 00:25:15 | 00:30:18 | 83.33 | 00:05:03 | 00:00:00 |
| yBYI | 00:10:06 | 00:10:06 | 100 | 00:00:00 | 00:00:00 |
| yBYI | 00:10:06 | 00:25:15 | 40 | 00:10:06 | 00:00:00 |
| *Mean (yBYI)* | *00:12:38* | *00:23:59* | *60.02* | *00:06:18* | *00:03:47* |
| oFZY | 00:20:12 | 02:51:38 | 11.77 | 00:55:32 | 00:05:03 |
| oFZY | 01:30:51 | 05:23:00 | 28.13 | 00:10:06 | 00:00:00 |
| oFZY | 00:15:09 | 02:56:38 | 8.58 | 02:11:12 | 00:30:17 |
| oFZY | 00:40:22 | 03:32:00 | 19.04 | 00:10:05 | 01:15:43 |
| oFZY | 01:05:38 | 05:12:59 | 20.07 | 00:30:18 | 00:05:03 |
| oFZY | 00:05:03 | 01:20:46 | 6.25 | 00:15:09 | 01:00:34 |
| *Mean (oFZY)* | *00:39:32* | *03:32:50* | *15.79* | *00:42:04* | *00:29:27* |
| gZUN | 00:35:00 | 00:40:00 | 87.50 | 00:00:00 | 00:05:00 |
| oFIJ | 01:10:41 | 01:40:56 | 70.03 | 00:05:02 | 00:15:08 |
| *oFIA | *N/A* | *N/A* | *N/A* | *N/A* | *N/A* |

**Table S4:** Percentage of female nest recesses with males’ nest sitting during incubation. We give the number of recesses females took and the number of nest sitting events of the males as well as the number of days during which the wildlife cameras were functioning. Two individuals, denoted in grey, were excluded when calculating percentages. For male gZUN the camera worked reliably for one day only, and male oFIA was excluded because he sat on the nest only after eggs hatched.

| **Nest sitting males** | **N female recesses** | **N males nest sitting** | **% nest recesses with male nest sitting** | **N days when cameras were functioning** |
| --- | --- | --- | --- | --- |
| oFYJ (201913) | 12 | 4 | 33.33 | 3 |
| yBYI *(mate yAUF w/transmitter 201919)* | 98 | 4 | 4.08 | 13 |
| oFZY (201918) | 14 | 6 | 42.86 | 4 |
| gZUN (201915) | 1 | 1 | 100 | 1 |
| oFIJ (201924) | 98 | 1 | 1.02 | 15 |
| *oFIA (201930) | N/A | N/A | N/A | 19 |

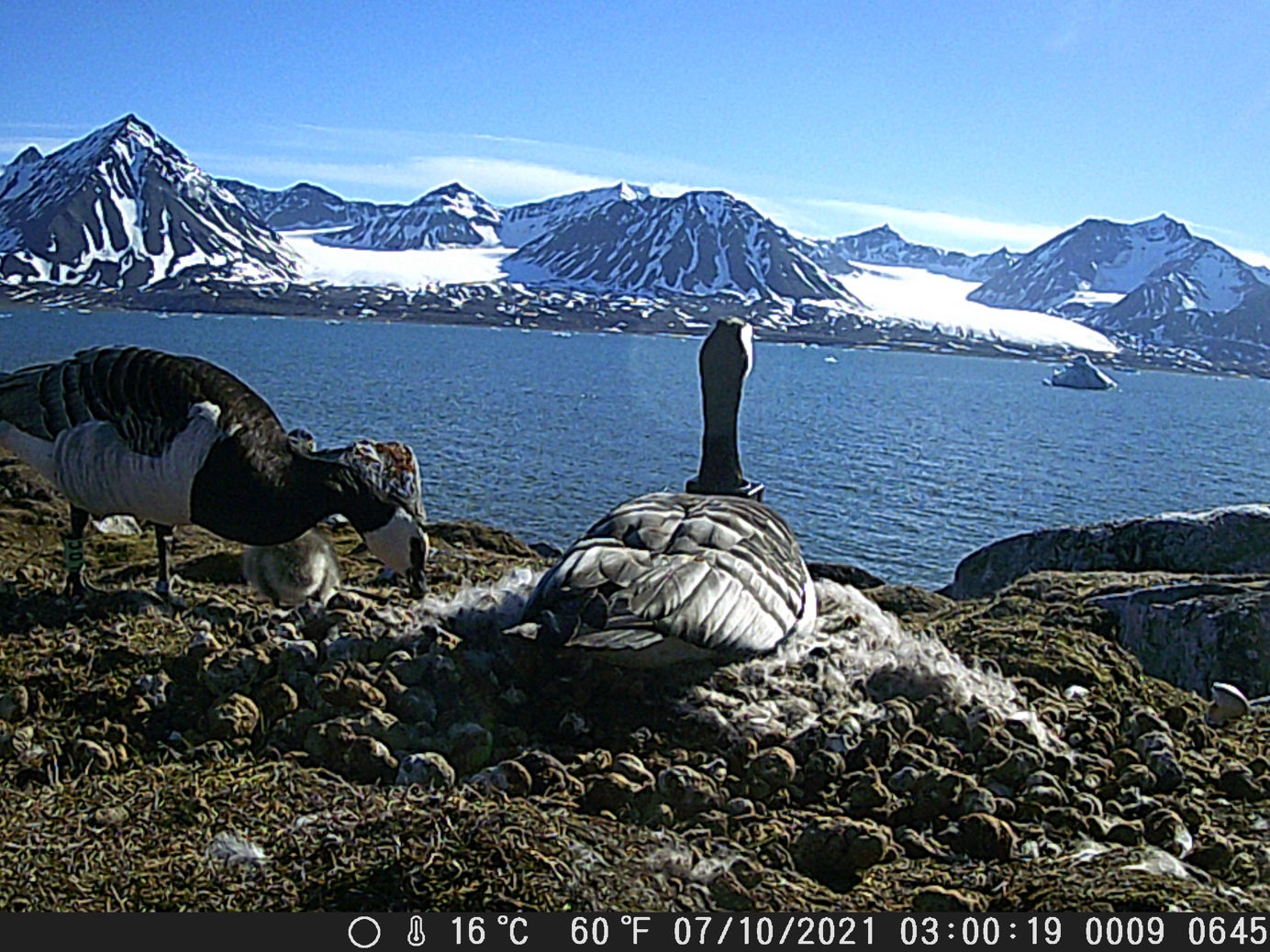

**Figure S1:** Male oFIA began nest sitting shortly after all eggs hatched. His mate gLDC stayed in close vicinity to the nest with the newly hatched three goslings. (Photo taken from the wildlife camera. © M. de Jong & I. B. R. Scheiber)
